# A Basal Ganglia Model for understanding Working Memory Functions in Healthy and Parkinson’s Conditions

**DOI:** 10.1101/2023.07.04.547640

**Authors:** Vigneswaran C, Sandeep Sathyanandan Nair, V. Srinivasa Chakravarthy

## Abstract

Working memory is considered as the scratchpad to write, read, and process information to perform cognitive tasks. Basal Ganglia (BG) and Prefrontal Cortex are two important parts of the brain that are involved in working memory functions and both the structures receive projections from dopaminergic nuclei. In this modelling study, we specifically focus on modelling the working memory functions of the BG, the working memory deficits in Parkinson’s disease conditions, and the impact of dopamine deficiency on different kinds of working memory functions. Though there are many experimental and modelling studies of working memory properties, there is a paucity of models of the BG that provide insights into the contributions of the BG in working memory functions. The proposed model of the BG is a unified model that can explain the working memory functions of the BG over a wide variety of tasks in normal and Parkinson’s disease conditions.

## 1. INTRODUCTION

Basal Ganglia (BG) is long known for its critical role in Working Memory (WM) along with its attributions to Reinforcement Learning (RL) and motor action selection (Goldman-Rakic, 1995; McNab & Klingberg, 2008). There are many observations of high activity in BG and Prefrontal Cortex (PFC) during WM tasks, but the BG and dopaminergic projections from BG to PFC are considered to perform gating of items and saliency computation to execute WM tasks (Braver & Cohen, 1999; Dipoppa et al., 2016; Lisman et al., 1998; Volle et al., 2005). In attractor-based models of working memory, gating plays a key role in state transitions, persistence of memory and enhancement of stability against noise (Baddeley, 2010; Gonon, 1997; O. Gruber & von Cramon, 2003). Gating is thought to be implemented in BG by dopamine projections to striatal medium spiny neurons whose bistable dynamics are modulated by dopamine (Kawagoe et al., 2004; Nakahara et al., 2004). Thus, the dopamine gating in BG essentially attenuates less-salient noise and latches salient items to perform WM functions like storage, updation and manipulation of memory items (Kawagoe et al., 1998; MINK, 1996). Many studies demonstrate WM deficits in animal models with loss of dopamine neurons and report improvement in performance after delivery of dopamine agonist drugs (Miyoshi et al., 2002; Rusu et al., 2013). In Parkinson’s disease (PD), depletion of dopaminergic cells in Substantia Nigra pars Compacta (SNc) affects the patient in different motor modalities including tremor, rigidity, and bradykinesia in arm movements, micrographia in handwriting generation and, dysarthria in speech (Fekete & Jankovic, 2011; Magdoom et al., 2011; Mandali et al., 2016; Nair et al., 2022; Oliveira et al., 1997).

PD patients, like patients with frontal lobe lesions, exhibit significant impairments in terms of executive cognitive functions with attentional control deficits (Engle, 2002; Gick et al., 1988). Cognitive tasks involving medial temporal-lobe structures are shown to be less impacted among PD patients (Owen et al., 1992). The ‘frontal-like’ Working Memory (WM) tasks are considered as the benchmark to analyse the severity of the disease and a discriminant tool to analyse subgroups of PD patients (Owen et al., 1993). However, some studies have shown that decrease of performance in spatial WM tasks is more pronounced than in non-spatial WM tasks, among moderate disease level patients (Owen et al., 1993; Takahashi & Hatakeyama, 2011). There are many experiments in literature that show WM deficits among PD patients such as, backward digit span and ordering task (Bublak et al., 2002) and, random number generation task (Robertson et al., 1996). Studies show that the difference in storage capacities between control and PD patients are not significant but the difference is more prominent in executive functions (Cooper et al., 1991; Dalrymple-Alford et al., 1994). Executive function impairments of WM are found to be very specific and it is still unknown why some WM functions are intact among PD patients but others are affected (Belleville et al., 2003; Miyake et al., 2000; Owen, 2000). (Gabrieli et al., 1996) suggested that reduced psychomotor speed observed among PD patients indicates the inability to perform complex cognitive functions that are necessary for performing executive WM tasks.

Computational and mathematical models at various levels of abstraction from detailed, biophysical single neuron to abstract networks of neuron-like units have been proposed to understand the underlying mechanism of BG and dopamine’s influence on WM (Balasubramani et al., 2015; Frank et al., 2001; Gruber et al., 2006; Nair et al., 2022; O’Reilly & Frank, 2006). Using a conductance-based neuronal network model of prefrontal cortex, (Durstewitz & Seamans, 2002) showed that dopamine’s modulation of the network via D1-receptors, resulted in deepening and widening of the attractor basins of WM states. The gating property of dopamine in WM is also modelled using spiking neurons (Brunel & Wang, 2001). Dopamine precursors like Levodopa administration to PD patients have shown improvement in various kinds of WM tasks (Floel et al., 2008; Simioni et al., 2017).

In more abstract versions, models showing the gating mechanism of dopamine on striatum are proposed without the biological details of gating (Dreher & Grafman, 2002; O’Reilly et al., 1999). There are also more abstract, systems-level models to replicate the key mechanisms of BG in WM (Frank et al., 2001; Gruber et al., 2006; Nakahara et al., 2004; Schroll et al., 2012; Trutti et al., 2021). However, abstract models also have limitations such as inability to show the relation between genetic/molecular level and systems level, and the inability to generalise to novel tasks (Gilbert et al., 2005; Lewis, Dove, et al., 2003; Schroll & Hamker, 2013). These limitations hinder the understanding of trends observed by the same subject in different experimental tasks and functions, and impedes us to understand the characteristic outcome of neurodegenerative diseases. Experimental data from different cognitive tasks performed by PD patients, under medicated and unmedicated conditions, show specific WM impairments and the reasons for which are still not clearly known, when compared to other modalities like motor, speech, and gait & balance (Cai et al., 2022; Ye et al., 2022). There are only a handful of BG models that incorporate the PD condition in WM tasks; and even they are often not general enough to describe PD-related WM deficits in a variety of WM tasks described in the literature (Goulding et al., 2022; Hazy et al., 2007; Schroll et al., 2012).

We propose a generalised model of BG under both control and PD conditions, and use it to explain the experimentally reported results of different WM tasks. The proposed model is validated in terms of its ability to show the specific executive WM impairments observed among PD patients. The model is validated against different WM tasks and in some tasks, the model performance is compared against experimental data reported in the original studies for both control and PD conditions. The model is also showcased to predict response times in different WM tasks. The subsequent section titled Methods and Materials describes the various tasks modelled to assess the WM functions of the BG model proposed. This is followed by the Results section where performance of the model in both control and PD conditions on the tasks described in the previous section is described. In the Results section, we also showcase the prediction capacity of the model in terms of model response times for tasks that do not have experimental response time and levodopa study on PD model to present its effects on WM performance. We then conclude by discussing the results, limitations, and the future scope in Section 4.

## 2. MATERIALS AND METHODS

### 2.1. Experimental Tasks

The proposed BG model can simulate a variety of experimental paradigms that can test sequence learning and working memory functions. Behavioural research uses a variety of tasks to assess sequence processing and working memory capabilities in humans. Sequence processing often implies working memory capability, since many sequence processing tasks require the sequence information to be stored for a finite duration, in order to produce the output sequence. Therefore, in this study we only use sequence processing tasks that involve working memory functions. Taking these aspects into consideration we modelled five WM tasks (**Figure 1**.). Of the five tasks, 1AX2BY task is chosen to test the sequence processing capacity of the model; Digit span task to validate the generalisation of the model in standard WM function; and Alphabetical Manipulation, Updating and 4-Consonants tasks are selected to check whether the PD version of the model can show specific impairment in executive WM function but intact manipulation of WM function. For all the tasks, Mean Squared Error (MSE) and Adam are used as cost function and optimizer respectively. All the implementations are done using Tensorflow 2 library.

**Figure 1.**
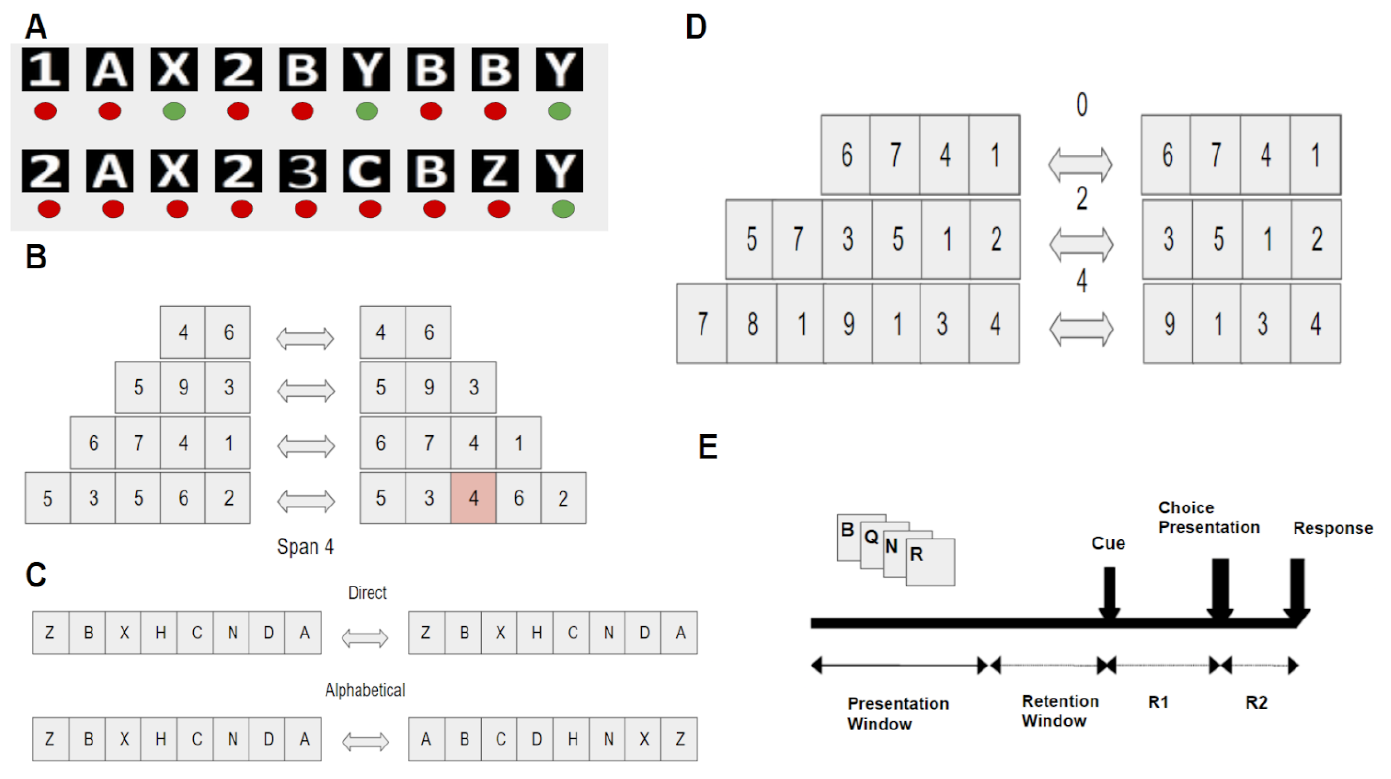
Schematic explanation of different working memory tasks modelled in this study. (A) 1AX2BY task - when the string AX (BY) follows 1(2) then the “Right” option should be chosen. (B) Digit span task - a sequence of digits should be recalled and the maximum spanning capacity of the model is calculated. (C) Alphabetical manipulation task - in “direct” case, the same sequence of letters presented to be recalled and in “alphabetical” case, the sequence should be recalled in dictionary order. (D) Updating task - recall the last ‘k’ number (k-span capacity) of items with different sequence lengths containing ‘n’ extra items (n-number of updates). (E) 4 consonants task - recall a sequence of 4 consonants based on the instruction given during instruction cue time step.

a. 1AX2BY task: The task involves sequences that comprise two types of stimuli - numeric stimuli and alphabetical stimuli (Frank et al., 2001). When the numeric 1 is encountered first in the input sequence, action ‘Right’ is the desired action only when A followed by an X also appears in the sequence. For all other items in the sequence, action ‘Left’ is the desired action. Similarly, when the numeric 2 is encountered first, action ‘Right’ is the desired action only when B followed by Y appears in the sequence. Similarly, for all other alphabetical sequences, action ‘Left’ is the desired action. Along with these items, in the sequence three noise input stimuli (3, C & Z) are also presented, which should be ignored.
b. Digit Span task: In case of the Digit Span task, the subjects are presented with a sequence of numbers (Gilbert et al., 2005). The length of the sequence is incrementally changed starting from two till the maximum length that can be successfully recollected. The length of the sequence that can be reliably recalled by the person is considered as the span of that person. The span of the model is evaluated by a procedure similar to the one described in (Gilbert et al., 2005). Starting with sequence length of 2, the model is expected to reproduce at least two sequences in six trials to move to length(sequence)+1. If the model fails to reproduce at least two sequences of six trials then the previous sequence length is taken as the span capacity.
c. Alphabetical Manipulation task: In case of the alphabetical recall task, the subject is presented with a sequence of alphabets (Gilbert et al., 2005). There are two variations of this task. In the first case (direct), the sequence presented is to be recalled in the same order. In the second case (alphabetical) the sequence of alphabets presented is to be recalled in the sorted order (A-Z).
d. Updating task: In this task, the subject is presented with a sequence of alphabets and must recollect the last k-items (Gilbert et al., 2005). Usually, ‘s’ (number of span items) is chosen to be the same as the span of the subject and appended with ‘n’-extra items (n-number of updates). In this task, the difficulty depends on the number of items added to the baseline sequence as it is commensurate with the number of updates required in the memory. The model is trained using sequence length of *s* along with *n* items, and however during retrieval the model is trained to recall the last *s* number of items. For example, when the sequence of length *s+n* is given as input, the model should recall last *s* items.
e. 4-Consonants Task (BQNR Task): Here three scenarios are considered. In the first case (same), the subject must recall the sequence in the same order as presented (Lewis, Cools, et al., 2003). In the second case (ends), the subject should recall based on the (3412) rule (for example, the input sequence BQNR is to be retrieved as NRBQ). In the third case (middle), the middle elements should be swapped while recalling (for example sequence BQNR is retrieved as BNQR). For modelling, four consonants are randomly chosen from seven consonants for each trial and each consonant is encoded to a 7 bit on-hot encoder. The sequence has four parts: i) 4 items are input consonants, ii) d items of empty input (7-bit vector with all zeros), iii) 1 item of instruction cue to mention the type of manipulation to be performed and followed by iv) 2 items representing two choices among which one must be selected.

### 2.2. The Model description

We present a model of the BG that can simulate the experimental tasks mentioned above **Figure 2**. (Nair et al., 2023). In this model, the input subsystem, Striatum, receives cortico-striatal projections from the cortical neurons, and the output nucleus - Globus Pallidus interna (GPi) produces the output sequence. The Subthalamic nucleus (STN) and the globus pallidus externa (GPe) together constitute the Indirect Pathway, whereas the direct striatal projections to GPi, constitute the Direct Pathway.

**Figure 2.**
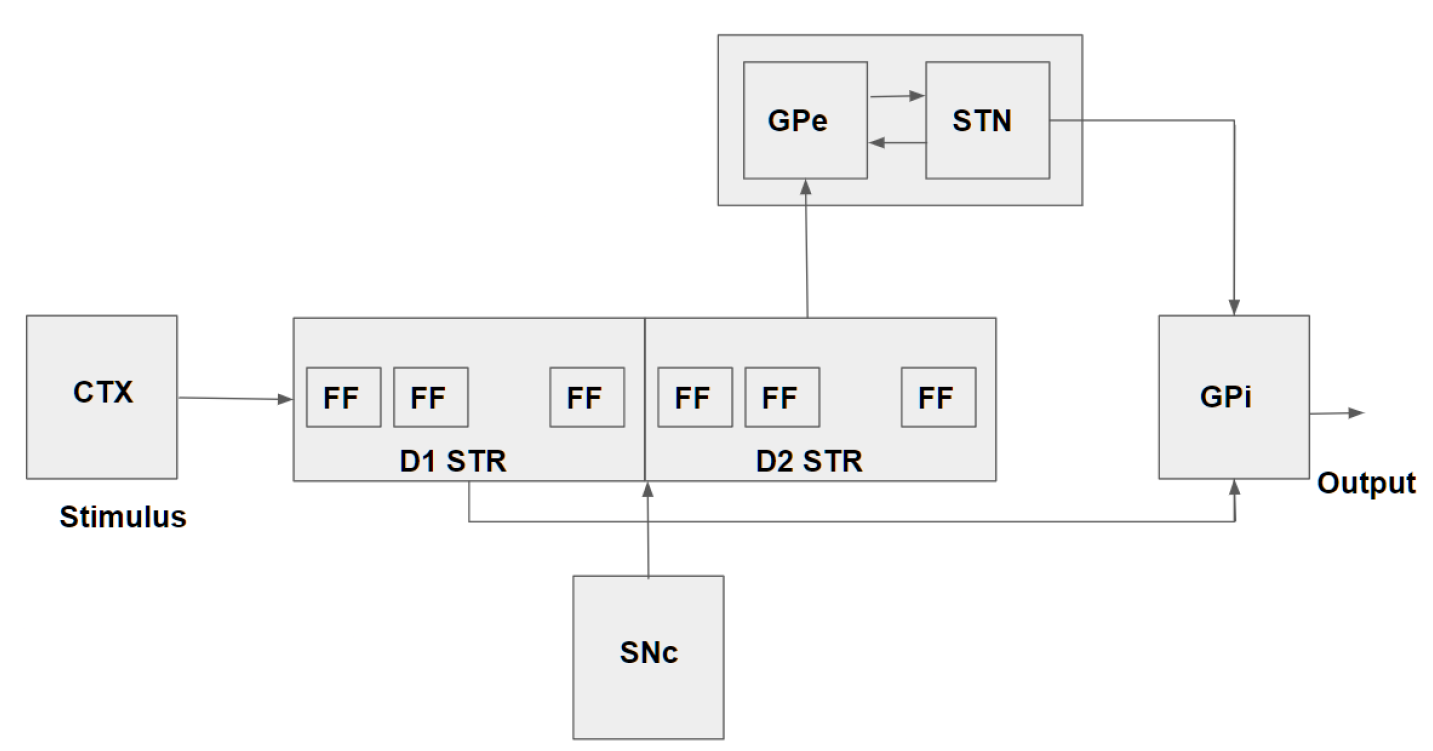
Block diagram of the proposed BG model. Input from the cortex (CTX) is presented to the Striatum (STR) - the input port of Basal Ganglia (BG); Striatum consists of D1 and D2 neurons modelled using Flip Flop (FF) neurons. The STR projects to the output port, Globus pallidus interna (GPi) via the direct pathway and indirectly via the indirect pathway. The Globus pallidus externa (GPe) and the subthalamic nucleus (STN) subsystems constitute the indirect pathway. Substantia Nigra pars compacta (SNc) consists of dopamine producing neurons, which regulate the information flow through the direct and indirect pathways.

#### 2.2.1. The Cortico-Striatal Projections

Cortical projections to the BG are received at the input nucleus Striatum. The striatum consists of two blocks labelled: D1 and D2. The D1(D2) - striatum consists of neurons that represent D1(D2)-receptors expressing Medium Spiny Neurons (MSNs). The MSNs are known to exhibit up/down states that are thought to underlie working memory functions (Plenz & Kitai, 1998). In the present model, the up/down dynamics of the neurons of D1- and D2-striatum are modelled using flip-flop neurons (Kumari et al., 2023). The flip-flop neurons, that are fashioned after the flip-flops of digital systems, have proven memory capabilities (Kumari et al., 2023). The flip-flop neurons have two inputs (J and K) and a single output (V). The dynamics of the flip-flop neurons are given by eqns. (1-8) below.

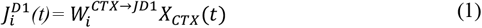

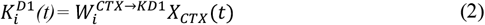

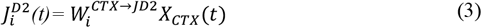

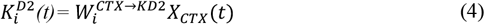

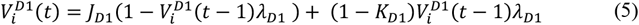

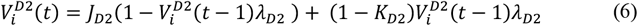

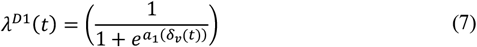

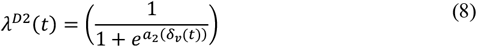

where *a*_1_ < 0 and *a*_2_ >0 and are the sigmoid gain parameters, 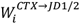 represent the weights connecting cortex and striatum, *X*_*CTX*_(*t*) denotes the input to cortex at time ‘t’, λ^*D*1/2^ is the dopamine gating variable and Value function (VF) is as given by Eqn. 9 below.

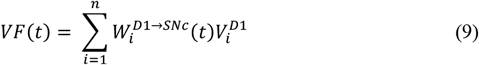

where 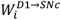 is the weight projecting from SNc to D1 striatum. Since D1- and D2-MSNs are modulated by dopamine, the flip-flop neurons are modulated by dopamine via the slope variables D1 and D2 as given by equations (7 and 8). The quantity *δ*_*v*_(*t*) is given as,

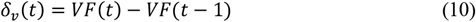

#### 2.2.2. The Output, GPi

GPi is the output of the BG and receives inputs from the striatum via both the direct and the indirect pathways. In the case of direct pathway, the D1 striatal neurons directly project to the GPi. The weights 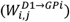 between the D1 MSNs and the GPi neurons are updated using Q Learning eqns. (22 and 23). The STN-GPe subsystems constitute the indirect pathway and are inter-connected via excitatory and inhibitory projections, respectively. Thus, STN-GPe system forms an excitatory-inhibitory loop and has been shown to exhibit different dynamical regimes. It has been thought to act as an exploration engine by adding a controllable level of noise to GPi (Chakravarthy & Balasubramani, 2014). STN-GPe dynamics are modelled by the following equations,

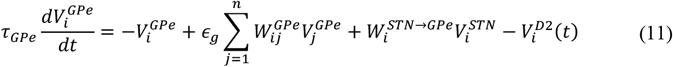

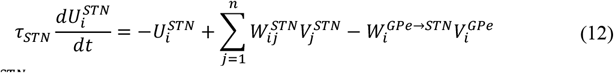

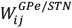 lateral weights within GPe/STN are initialised by,

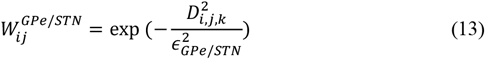

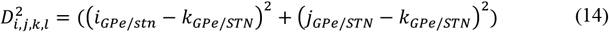

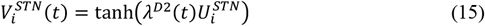

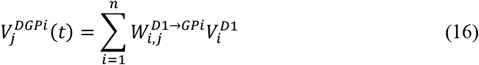

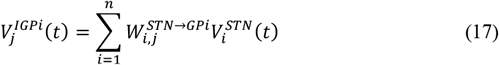

where *i*_*Gpe*/*stn*_, *j*_*Gpe*/*stn*_ and *k*_*Gpe*/*stn*_ are neuron indices to calculate the distance 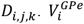 and 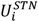 represent the internal states of GPe and STN neurons respectively, 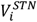 represents the output of the STN neurons, *τ*_*Gpe*_ and *τ*_*STN*_ are the time constants of GPe and STN, respectively and, 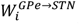 is the connection strength from GPe to STN, 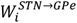 is the connection strength from STN to GPe,. 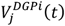 and 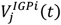 are the inputs arriving at GPi via direct and indirect pathways.

GPi neurons combine the outputs of D1-striatum and STN, and generate the predicted output as follows.

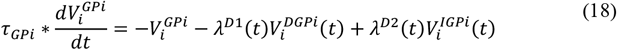

The GPi output (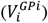) is compared against the threshold, *V*_*th*_, as a race model. Race models are used in modelling action selection in behavioural tasks, where *i*^*th*^ neuron whose output crosses the threshold first, is considered the winner, and the corresponding *i*^*th*^ action is selected (Kumari & Chakravarthy, 2022). The number of steps taken to cross a predefined threshold is equivalent to the response time of the model.

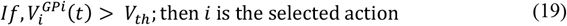

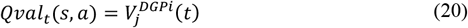

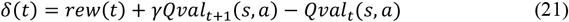

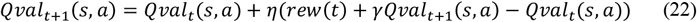

where, *γ* is the discount factor and 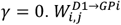 is updated using.

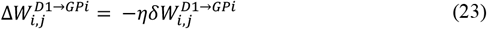

#### 2.2.2. The PD condition

In order to simulate the PD condition of dopamine deficiency, we clamp the dopamine variable, *δ*(*t*) to *δ*_*lim*_ in eqns. (7 and 8). If PD=1 then,

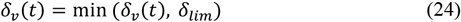

If PD_ON=1 (administration of Levodopa) then,

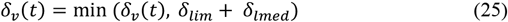

#### 2.2.3. Parameter selection

The list of parameters used in the model and their corresponding values are shown in **Table 1**. For three parameters ε_*Gpe*/*STN*_, *V*_*th*_and *δ*_*lim*_, we perform a parameter selection method with the suitable criterion for each of the parameters and use the obtained values for all simulations in this work. **Table** 2 shows the criteria for choosing suitable values for the three parameters.

**Table 1.**
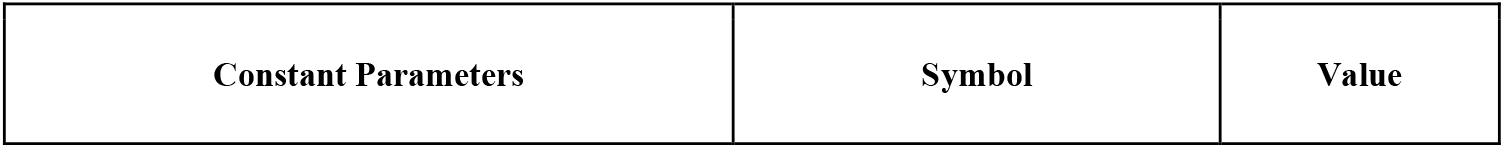
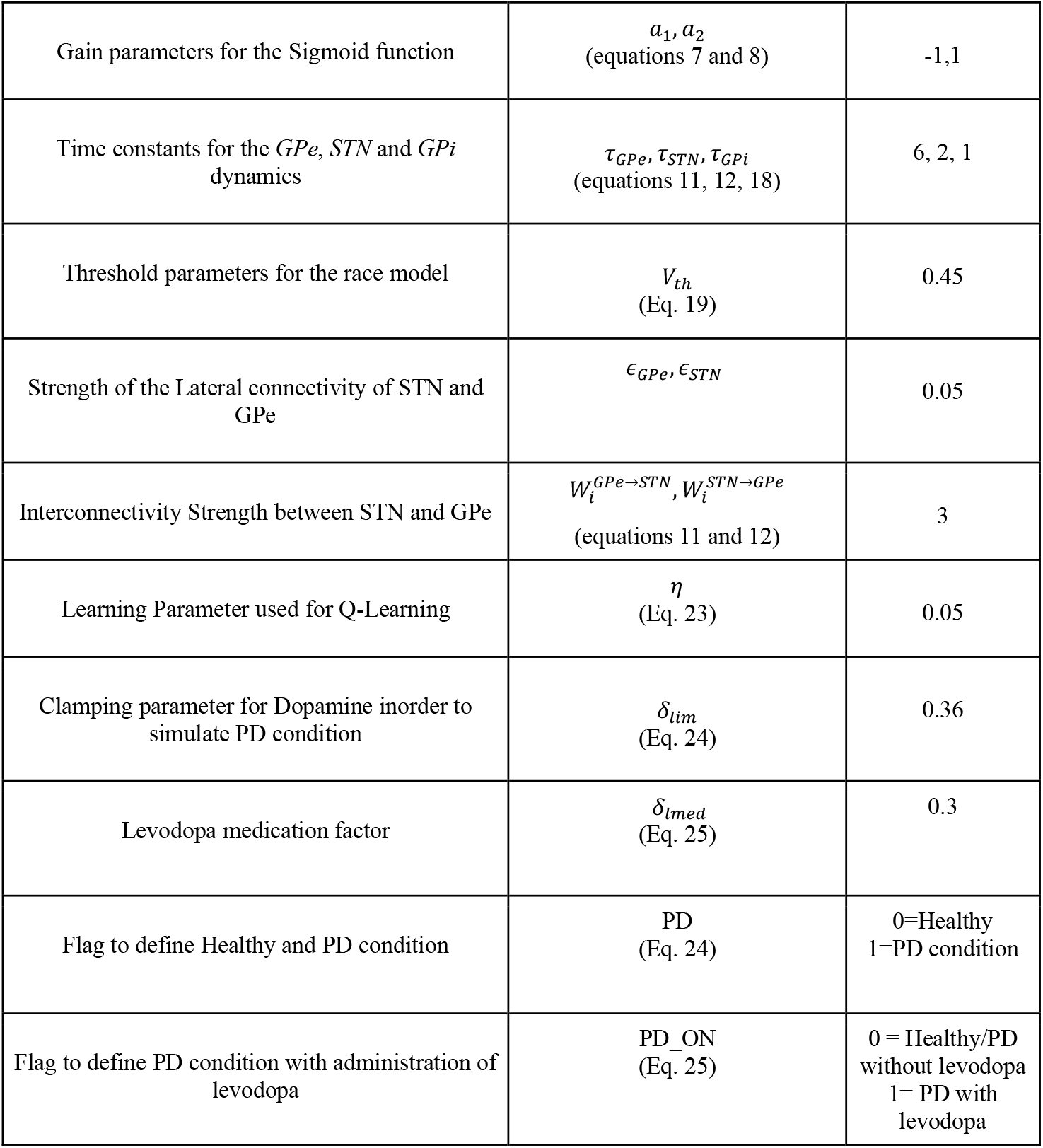
Model parameters

| Constant Parameters | Symbol | Value |
| --- | --- | --- |
| Gain parameters for the Sigmoid function | $a_1, a_2$<br>(equations 7 and 8) | -1,1 |
| Time constants for the *GPe*, *STN* and *GPI* dynamics | $\tau_{GPe}, \tau_{STN}, \tau_{GPI}$<br>(equations 11, 12, 18) | 6, 2, 1 |
| Threshold parameters for the race model | $V_{th}$<br>(Eq. 19) | 0.45 |
| Strength of the Lateral connectivity of STN and GPe | $\epsilon_{GPe}, \epsilon_{STN}$ | 0.05 |
| Interconnectivity Strength between STN and GPe | $W_i^{GPe \rightarrow STN}, W_i^{STN \rightarrow GPe}$<br>(equations 11 and 12) | 3 |
| Learning Parameter used for Q-Learning | $\eta$<br>(Eq. 23) | 0.05 |
| Clamping parameter for Dopamine inorder to simulate PD condition | $\delta_{lim}$<br>(Eq. 24) | 0.36 |
| Levodopa medication factor | $\delta_{lmed}$<br>(Eq. 25) | 0.3 |
| Flag to define Healthy and PD condition | PD<br>(Eq. 24) | 0=Healthy<br>1=PD condition |
| Flag to define PD condition with administration of levodopa | PD_ON<br>(Eq. 25) | 0 = Healthy/PD<br>without levodopa<br>1= PD with<br>levodopa |

**Table 2.** Parameters and their selection criterion

| Parameter | Criterion |
| --- | --- |
| $\epsilon_{GPe/STN}$ | Maximum standard deviation of noise from STN-GPe |
| $V_{th}$ | Closely matching response time from experimental data of 4-consonants (same) task in control condition |
| $\delta_{lim}$ | Closely matching accuracy from experimental data of 4-consonants (middle) task in PD condition |

*∈*_*Gpe*/*STN*_ controls the variance in noise from STN-GPe and acts as an exploration unit for learning or performing a task. Thus, the *∈*_*Gpe*/*STN*_ value is chosen to generate the maximum noise variance. *V*_*th*_ controls the GPi race model iterations and thus the response time of the model. Small *V*_*th*_ may not be sufficient to give accurate predictions whereas large *V*_*th*_will unnecessarily make the race model loop to run for the maximum number of iterations. Therefore, experimental data of control response time from 4-consonants task (same - condition) is used as the criterion for choosing *V*_*th*_. *δ*_*lim*_ controls the clamping of dopamine and therefore, the suitable value is obtained from the matching of PD experimental data accuracy of 4- consonants task (middle - condition). The values obtained from the parameter selection process is used in all the experiments. **Figure 3**. shows the plots of the cost function for corresponding parameter values.

**Figure 3.**
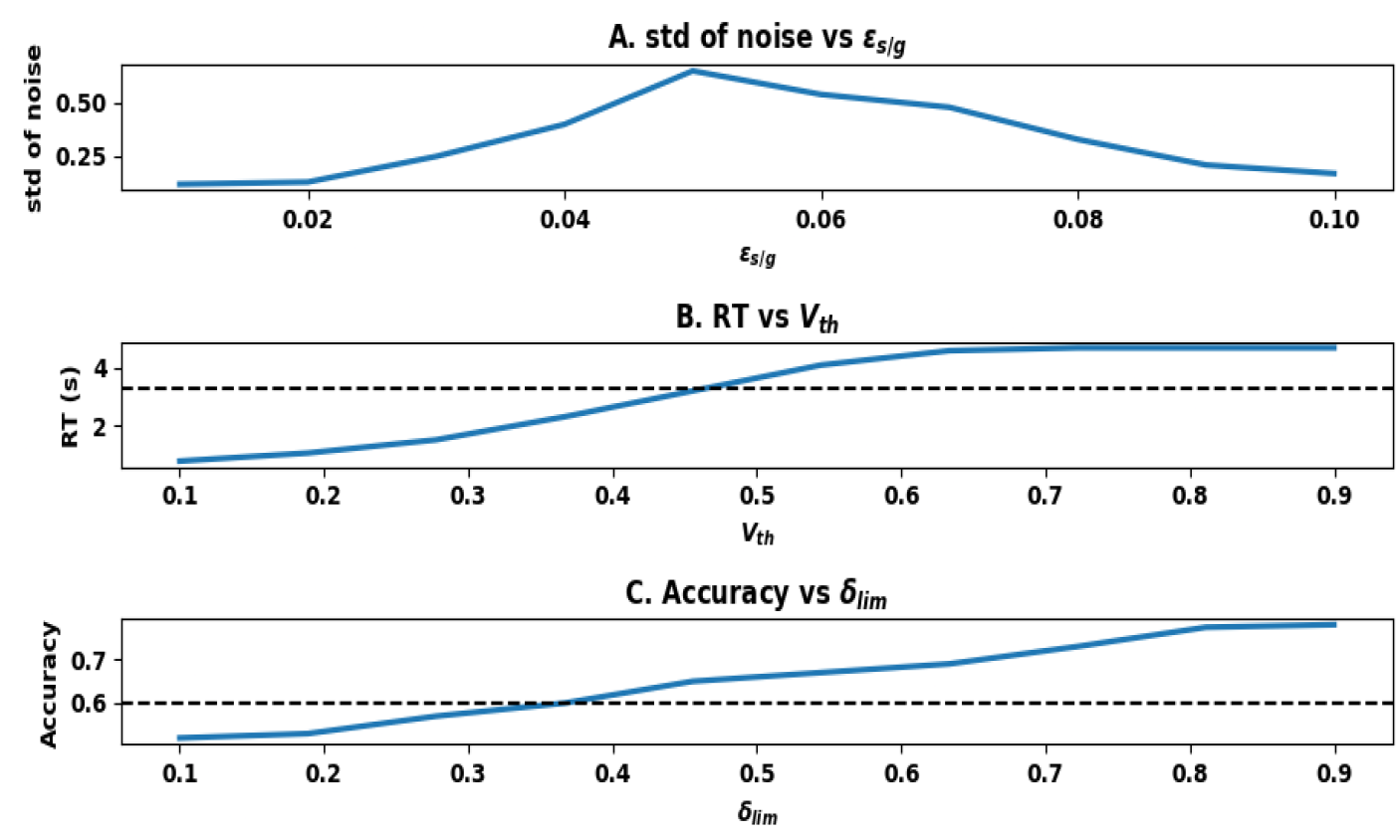
(A) Standard deviation of noise from STN-GPe as a function of the lateral weights of STN-GPe *∈*_*Gpe*/*STN*_. (B) Response time in seconds when varying *V*_*th*_. Black dashed line marks the response time given in the experimental data of the control subject in performing 4-consonants task (same - condition). (C) Accuracy of 4-consonants task (middle - condition) when varying *δ*_*lim*_. Black dashed line marks the accuracy of PD subject provided in experimental data.

**Table** 3 shows the network architecture sizes for different tasks. The architecture sizes are selected based on task requirements.

**Table 3.** Network parameters for different tasks

| Task | No. of Striatum D1/D2 neurons | No. of STN-GPe neurons | No. of GPi neurons |
| --- | --- | --- | --- |
| 1AX2BY | 20 | 20 | 2 |
| Digit span task | 200 | 50 | 9 |
| Alphabetical manipulation | 400 | 100 | 26 |
| Updating task | 400 | 100 | 26 |
| 4 - consonants task | 200 | 50 | 2 |

## 3. RESULTS

In order to validate the learning capacity of the proposed model, we show the performance in terms of accuracy and loss function for the 1AX2BY task. On extending the same, to check whether the proposed model can replicate the performance of PD subjects on WM tasks, we use four different tasks: digit span, alphabetical manipulation, update, and 4-consonants tasks, and compare model performance with experimental data for both control and PD conditions.

### 1AX2BY task

The model is trained for 50 epochs with a total number of 1860 data samples (batch size: 32, validation split: 0.3) and the sequence length of each sample is 20. **Figure 4** shows the accuracy and loss (mean and standard deviation) curves for both training and validation data with 10 repeated trials. The results indicate that the model can learn the task and generalise well.

**Figure 4.**
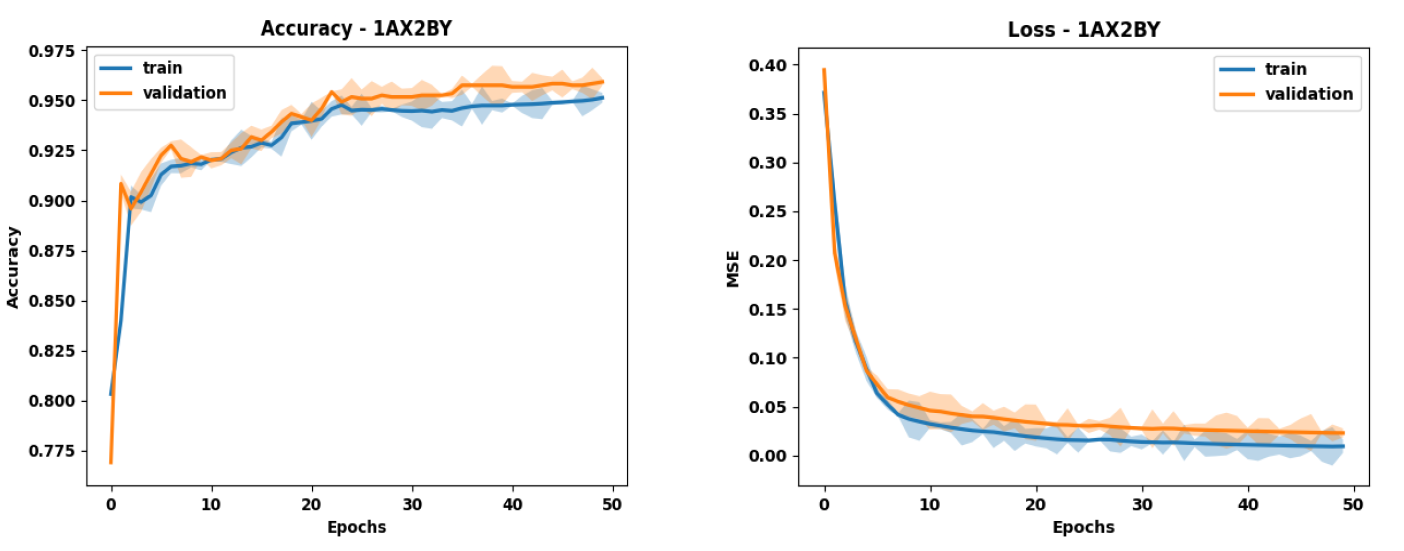
1AX2BY task - Accuracy and MSE loss curves of the model for train and validation data

### Digit span task

The standard digit span task is used to showcase the WM storage capacity of the model. The model is trained for 50 epochs for each sequence length, with 32 samples of the particular sequence length. The experiment is carried out in both control and PD conditions for 10 repeated trials. **Figure 5**. shows the span of the proposed model and experimental data as reported by (Gilbert et al., 2005). The results validate that in both control and PD versions of the model, there is no statistical difference (p > 0.05, n=10) between span of control and PD, and is consistent with the experimental observation.

**Figure 5.**
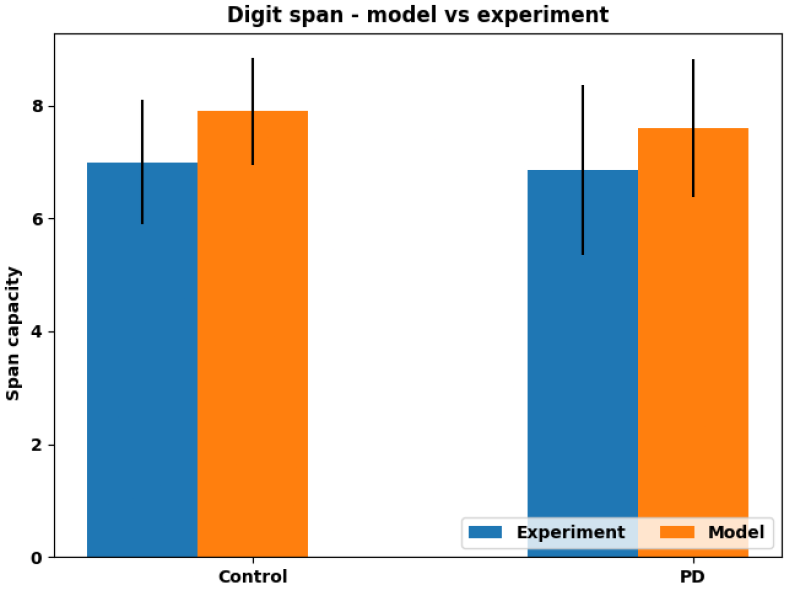
Spanning capacity of model compared with experimental data in both control and PD condition

### Alphabetical manipulation task

The alphabetical manipulation task is used to test whether the model can show a significant difference between control and PD in performing alphabetical recall task, whereas in direct recall, the performance is not significantly different between the two conditions. The span capacity of 8 obtained from the digit span task is set as the sequence length. The model is trained for 200 epochs, with English alphabets. **Figure 6** shows the accuracy of the proposed model and experimental data for direct and alphabetical recall as reported in (Gilbert et al., 2005) for both control and PD conditions. The significant difference observed in alphabetical recall task performance (p < 0.05, n=10) and non-significant difference (p > 0.05, n=10) in direct recall tasks are consistent with experimental observations.

**Figure 6.**
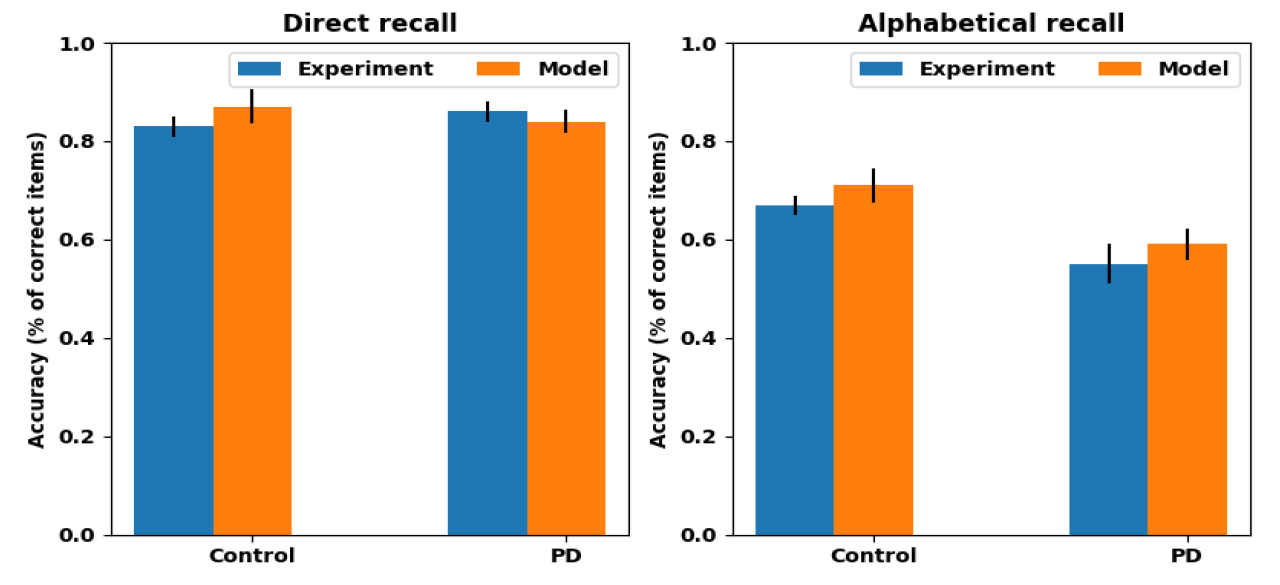
Comparison of model performance in terms of accuracy with experimental task in direct and alphabetical recall tasks for both control and PD conditions

### Updating task

Updating task requires updating of items in the stored list. The model is trained for 200 epochs with English letters, where each letter is transformed to a one-hot vector similar to the alphabetical manipulation task. **Figure 7**. shows the percentage of items recalled by the model and experimental data reported by (Gilbert et al., 2005) when k number of updates (k = 0, 2, 4, 6) are required for both control and PD conditions. The result shows the model performance is consistent with the experimental data and the differences between update groups is significant (p < 0.01).

**Figure 7.**
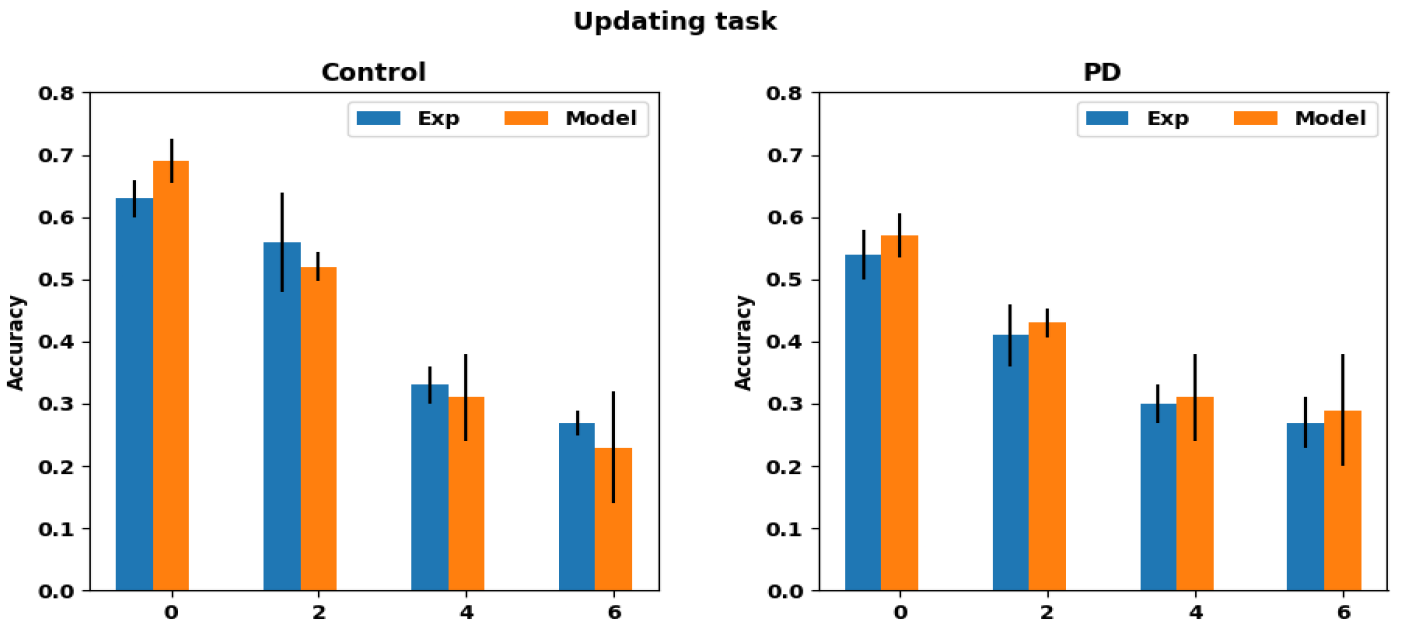
Accuracy of recalled items in updating task compared with model and experimental data for four different numbers of updates in both control and PD conditions

## 4 Consonants Task

The model performance on 4-consonants WM task in terms of accuracy and reaction time is compared against the experimental data reported by (Lewis, Cools, et al., 2003). In the experimental data, two parts of reaction times (R1 and R2) are reported. In order to compare with the model, both R1 and R2 in experimental data are added and compared. The response time of the model is equal to the number of steps taken by the winning GPi neuron to cross the threshold as in Eqn. 19. **Figure 8**. shows the model performance and experimental data for both control and PD conditions in terms of accuracy and reaction time (in seconds) grouped by different manipulation instructions. The results are consistent with experimental observations that all three manipulation groups are significantly different within each other (p < 0.01, n=10).

**Figure 8.**
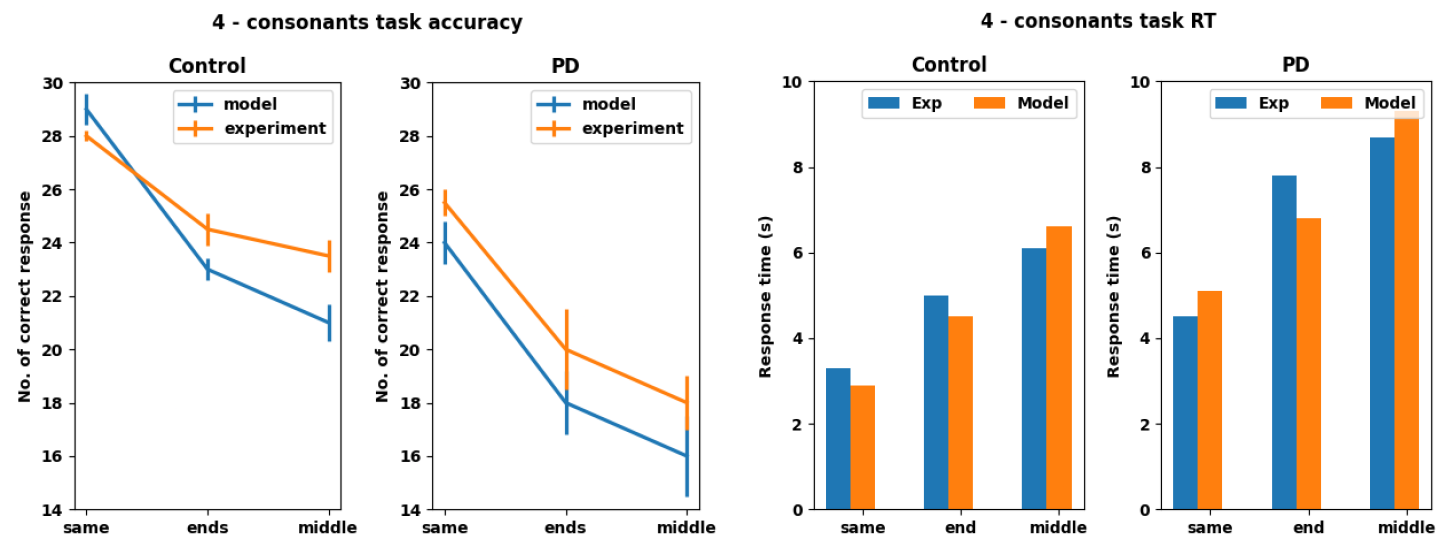
Comparison of model with experimental data for both control and PD conditions in terms of accuracy and reaction time

### Response time prediction

With the proposed model, we also predict the response time for digit span, alphabetical manipulation, and updating tasks. This shows that the model not only has the potential to match the experimental data but also can give the prediction of metrics missed in experimental data. Similar to the 4-consonants task, response time of the model is calculated based on the number of steps elapsed in the GPi race model, and the same experimental setup is used. **Figure 9** shows the predicted response time for the three tasks. It can be inferred that predicted response times are inversely proportional to the performance accuracy.

**Figure 9.**
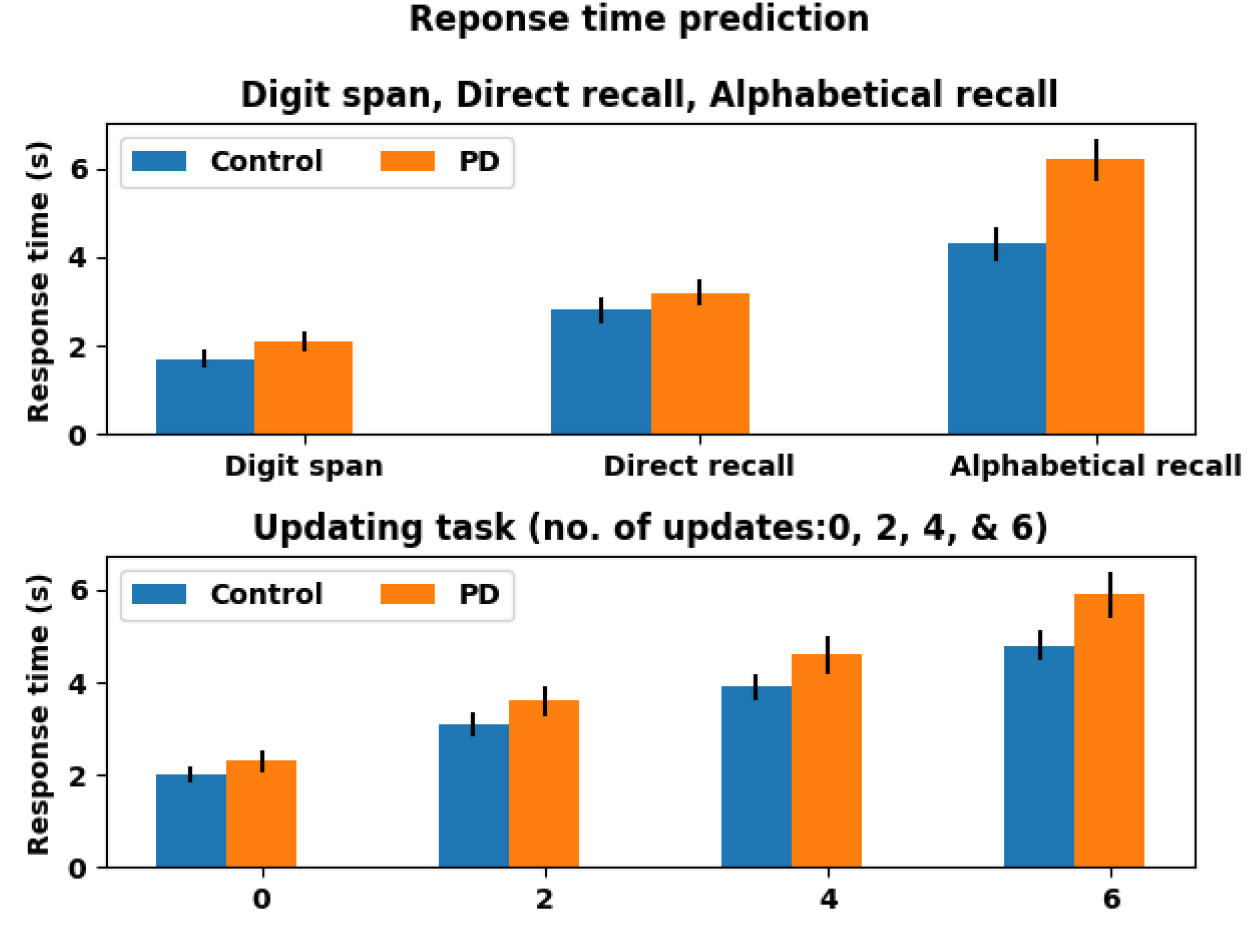
Comparison of predicted response time for digit span, direct recall, alphabetical recall, and updating tasks for both control and PD conditions

### Levodopa Effect

Using the proposed model, we study the effects of administering levodopa in 4-consonants task in terms of performance accuracy. **Figure 10** shows the performance of the model in three different conditions. It can be inferred the administration of levodopa improved the model performance in all three cases (same, ends and middle). Thus, model can be used as a potential test bench for testing the effect of administering medication to the patient in a clinical setting.

**Figure 10.**
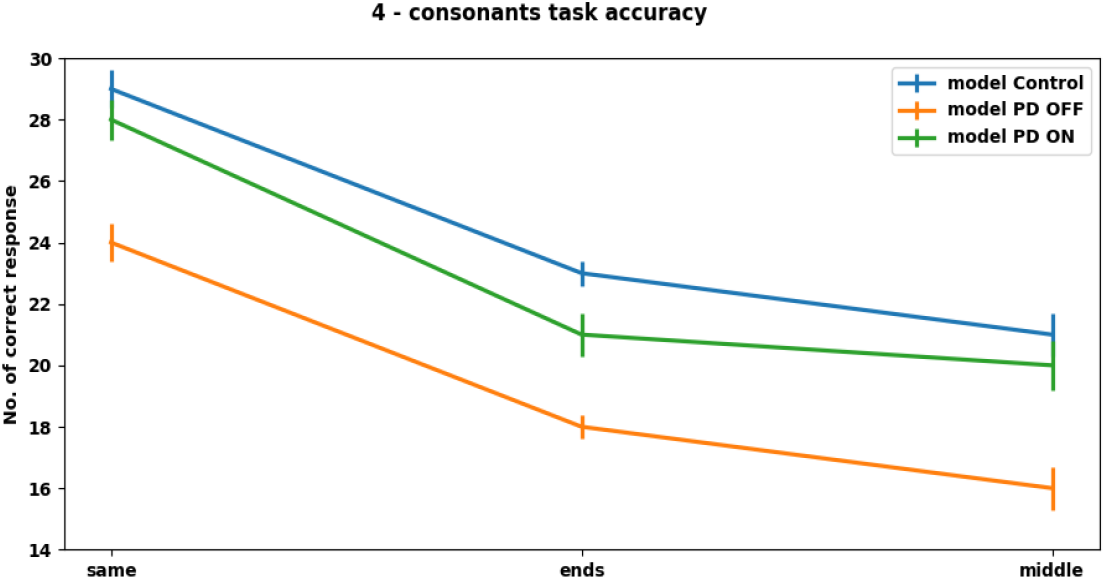
Effect of levodopa on PD model in 4-consonats task in terms of model performance accuracy. PD OFF and PD ON denote PD condition of the model without and with levodopa medications respectively

## 4. DISCUSSION

The proposed work presents a generalized basal ganglia (BG) architecture to model the WM functions of BG under normal and Parkinson’s conditions. The model architecture is cast in Reinforcement Learning (RL) framework and includes the memory feature implemented by the bistable flip-flop neurons in D1 and D2 Striatum, the input port of the BG; the output nucleus, Globus Pallidus interna (GPi) which implements action selection using a race model; the coupled system of non-linear oscillators that represent the STN-GPe oscillations in the indirect pathway, which was earlier proposed to supply exploratory drive necessary to implement RL. The proposed model could successfully implement the WM functions as well as capture the specific WM impairments observed among PD patients. The model can also predict response times of subjects in different WM tasks.

We chose five different tasks and the model results closely match with the experimental data reported in the original studies. The 1AX2BY task showed the learning capacity of the model on standard sequence processing and while the other four tasks Digit span, Alphabetical manipulation, Updating and 4-Consonants – were used to compare the performance in control and PD conditions. From our results, we could see that the digit spanning and direct recall tasks did not show any significant difference in the performance between the control and the PD cases, which is in line with the experimental results. However, there is a significant reduction in accuracy for the PD case for the alphabetical recall, updating and 4-consonant tasks and a significant increase in response time with respect to the 4-consonant tasks in PD. We could compare the response time only for the 4-consonant task due to the unavailability of response time for other tasks in experimental data. However, the model is used to predict the response time of tasks for which experimental data are unavailable. Overall, our model results closely resembled the trends in the experimental results.

From the wide range of literature that we discussed in relation to the role of BG in WM tasks we could infer that there is a lot of scope in exploring the working memory properties of BG and the related impairments caused due to PD. Computational and mathematical studies have explored dopamine’s influence and role of BG in WM functions (Balasubramani et al., 2015; Frank et al., 2001; Gruber & von Cramon, 2003; Nair et al., 2022; O’Reilly & Frank, 2006). There are also modelling studies representing the circuitry at more abstract levels and more detailed levels also (Braver & Cohen, 1999; Brunel & Wang, 2001; Dreher & Grafman, 2002). However, in this wide range of studies there is a conspicuous paucity of models that address alterations of WM functions under disease conditions such as PD. When it comes to cognitive aspects and WM deficits of PD there are only a few studies (Goulding et al., 2022; Hazy et al., 2007; Schroll & Hamker, 2013), and there is still a huge scope in studying different aspects of WM deficits in detail.

Our current work is a step in this direction. In this study, we have successfully modelled different types of WM with varying complexities. This helps us in comprehensively understanding working memory functions under different conditions and which aspects are impacted when the dopaminergic circuitry is disrupted. Currently, we only confine ourselves to the involvement of BG in WM and its impairments in PD, although WM involves different parts of the brain including PFC (Barbey et al., 2013). However, even within PFC it has been suggested that dorsolateral PFC supports spatial WM whereas ventrolateral PFC supports non-spatial WM (Barbey et al., 2013). In the current model, we only focus on non-spatial WM. Studies have shown evidence of WM in the posterior brain also, particularly in the parietal cortex (Curtis & D’Esposito, 2003). Thus, an integrative model of WM will necessarily include the task-relevant prefrontal, parietal structures in the cortex, and the BG in the subcortex. Creation of such an integrative model will define the goal of our future efforts in this area.

## DATA AVAILABILITY STATEMENT

The datasets generated during and/or analysed during the current study are available from the corresponding author on reasonable request.

